# Ancestry-linked stromal variations impact breast epithelial cell invasion

**DOI:** 10.1101/2024.12.26.630400

**Authors:** Madison Schmidtmann, Victoria Elliott, James Clancy, Harikrishna Nakshatri, Crislyn D’Souza-Schorey

## Abstract

Breast cancer is a significant health challenge worldwide, and disproportionately affects women of African ancestry (AA) who experience higher mortality rates relative to other racial/ethnic groups. Several studies have pointed to biological factors that affect breast cancer outcomes. A recently discovered stromal cell population that expresses <u>P</u>ROCR, <u>Z</u>EB1 and <u>P</u>DGFRα (PZP cells) was found to be enriched in normal healthy breast tissue from AA donors, and only in tumor adjacent tissues from donors of European ancestry (EA). Here, we investigated the effect of PZP cells on the acquisition of tumorigenic phenotypes in AA and EA human breast epithelial cells to determine its contribution to the biological basis of outcome disparities. Using 3D cell models of tumor invasion, we find that PZP cells confer invasive capacity to epithelial cells independent of genetic ancestry, wherein cells exhibit leader-follower behaviors during extracellular matrix invasion. Enhanced epithelial invasion stems from a combination of AKT activation and fibronectin deposition by the PZP cells. Although activation of AKT in epithelial cells alone is insufficient to induce invasive behaviors, blocking AKT activation markedly reduces invasive capacity. These findings point to the germaneness of differences in breast biology and the multi-faceted roles and enrichment in PZP cells in AA breast tissue, in furthering current understanding of the molecular basis for worse prognosis for patients of African descent.

## Introduction

Breast cancer is the most frequently diagnosed malignancy among women globally^1^. In the United States alone, this year an estimated 310,720 new cases of invasive breast cancer and 56,500 new cases of non-invasive (in situ) breast cancer will be diagnosed^2^. Another notable statistic is the observed decline in breast cancer mortality, largely because of early detection strategies and improved treatment options in adjuvant and metastatic settings. However, the mortality gap between women of African ancestry (AA) and white women of European ancestry (EA) persists^3^. Furthermore, population-based research finds that women of African descent have a higher incidence of aggressive and more invasive forms of breast cancer, such as triple negative breast cancer (TNBC), compared to women of other ancestries^4^. The disparity is even greater in the case of women under 50, where the mortality rate of AA women is double that of EA women and there is inherited predisposition to breast cancer among African American women^5^. Even when consequences of lower socioeconomic status and barriers to equitable care are accounted for, these outcome disparities remain, suggesting that biological factors may be an underlying driver^6–8^. In this regard, the African pan-genome contains ∼10% more DNA than the human reference genome of European origin^9^. Moreover, recent studies have included characterization of genetic ancestry, thereby deepening our understanding of the molecular mechanisms contributing to these outcome disparities^10,11^. The use of single nucleotide polymorphisms (SNPs) associated with geographical location has allowed researchers to stratify patient data based on genetic ancestry linked to continental location^12,13^. For example, it was found that tumor-associated immunomodulatory signatures are distinct for patients of African descent with TNBC. Similarly, by utilizing donor tissue defined by genetic ancestry, recent studies have identified a stromal cell population that expresses <u>P</u>ROCR, <u>Z</u>EB1 and <u>P</u>DGFRα (referred to as PZP cells) that are enriched in breast tissue of normal healthy AA donors but present in low abundance in healthy breast tissue from patients of other backgrounds^14^. Notably, the prevalence of PZP cells increases in tumor adjacent breast tissue of women with European ancestry^15^. The study also showed that PZP cells when transformed with oncogenes generate metaplastic carcinomas in NOD/SCID Gamma (NSG) mice pointing to their categorization as a cell-of-origin in metaplastic carcinomas of the breast.

Within a complex tumor microenvironment (TME), heterogenous populations of stromal cells are coopted for pro-tumorigenic and invasive roles^16,17^. For instance, fibroblasts are a plastic population of cells that under normal conditions function to control tissue homeostasis. During tumorigenesis, fibroblasts are converted to cancer associated fibroblasts (CAFs) which are known to influence cancer progression through secretion of factors such as IL-6, MMPs, and fibronectin, thereby enabling CAFs to affect the composition of the TME^18^. Intriguingly, stromal PZP cells harbor markers and characteristics that overlap with fibroadipogenic progenitors and mesenchymal stromal cells with the capacity to differentiate into either an adipogenic or myofibroblast lineages^15,19,20^. These cells also display CD105^high^/CD26^-/low^ markers that resemble interlobular human breast fibroblastic cells.

Given the properties of PZP cells, coupled with the differential enrichment in AA patients, we asked if PZP cells contribute to the acquisition of invasive phenotypes in human breast epithelial cells, supporting a biological basis for worse outcomes experienced by patients of African ancestry. We postulate that the intrinsically higher levels of PZP cells in normal breast AA tissues is a critical contributing factor to the accelerated onset and/or progression of breast cancer observed in AA women. Here, we report that PZP cells influence epithelial cell behavior through a combination of activating signals of cell invasion and the physical remodeling of the extracellular matrix. The PZP secretome promotes activation of AKT in epithelial cells of AA and EA genetic ancestries and blocking AKT activation significantly reduces epithelial cell invasion in a 3D spheroid co-culture model. However, activation of AKT alone is not sufficient to induce epithelial cell invasion and the presence of PZP cells is required to facilitate epithelial cell invasion. Moreover, PZP cells facilitate invasion through the combined action of proteolytic matrix degradation and fibronectin deposition. Together these results highlight multiple mechanisms through which PZP cells could influence the early stages of breast tumorigenesis and furthers our understanding of the molecular determinants that may contribute to the enhanced disease burden experienced by women of African ancestry.

## Results

### PZP cells confer invasive potential to normal epithelial cells

The properties of human KTB40 and KTB42 cell lines and other KTB cell lines utilized in this study are outlined in Table 1. KTB40 and KTB42 cell lines were propagated from the core biopsies derived from healthy breast tissue of women of African Ancestry (AA) donated to the Susan G. Komen Tissue Bank (KTB)^14^ that express <u>P</u>ROCR, <u>Z</u>EB1 and <u>P</u>DGFRα (hence named PZP cells) and are uniquely enriched in the normal breast tissue of women of AA descent^15^. Previously published genetic ancestry analysis of the PZP cell lines found that approximately 70% and 90% of SNPs examined in KTB40 and KTB42 respectively are of African ancestry^14^. To elucidate the impact of PZP cells on breast epithelial cell behavior, we utilized normal breast epithelial cell lines also derived from donated breast tissue to the Komen Tissue Bank^14^. As outlined in Table 1, cell lines KTB34 and KTB36 originated from EA populations and approximately 90% of genetic markers are of European ancestry. Cell lines KTB8 and KTB39, were derived from AA women and contain approximately 60% and 90% genetic markers of African heritage respectively^14^. To explore the influence of PZP cells on epithelial cell invasion, we employed a three-dimensional (3D) co-culture spheroid model that allows invasion to be monitored over time in a collagen I matrix. Each cell type was stained separately with a Cell Tracker dye to visually distinguish between the PZP and epithelial cells. Co-cultures of epithelial cells and PZP cells (1:1 ratios) or an equivalent number of epithelial cells alone were seeded in ultralow attachment conditions to form spheroids. Spheroids were transferred into collagen and imaged every 24 hours, and the distance traversed by epithelial cells through the collagen matrix was monitored. Monoculture spheroids of epithelial cells of both EA (KTB34 or KTB36; Fig. 1A, B. Suppl. 1A, B, C) and AA (KTB8 or KTB39; Fig. 1D, E. Suppl. 1D) origin, exhibited little to no movement into the collagen matrix. Monoculture spheroids of KTB40 (PZP) cells on the other hand, over time resulted in cell invasion radially into the matrix emanating from the center spheroid (Suppl. 1E). However, when epithelial cells were cocultured with PZP cells in 1:1 ratios, the resulting spheroids exhibited a significant increase in KTB epithelial cell invasion (Fig. 1A, B, D, E, Suppl. 1A, C, D). Both PZP cell lines, KTB40 and KTB42, behaved similarly in these assays and henceforth are referred to as PZP cells. KTB34 or KTB36 epithelial lines of EA origin when cocultured with PZP cells formed fully mixed heterogenous spheroids (Fig. 1A, Suppl. 1C). A similar phenotype was observed in KTB8 (of AA origin) and PZP cocultured spheroids (Suppl. 1D), whereas nearly 80% of KTB39 (of AA origin) and PZP cells organized into ‘lobes’ in cocultured spheroids, while the remaining spheroids exhibited more uniform mixing (Fig. 1D). Notably however, epithelial cells, regardless of ancestral origin, followed PZP cells which acted as leader cells^21^ during invasion into the matrix (Fig. 1C, F arrows, Suppl. 1F arrows). Epithelial cell movement was evident by 48 hours, and at longer time points. Additional PZP cells were observed moving into the collagen matrix at the edge of the spheroid. Quantification of epithelial invasive behavior revealed that greater than 80% of epithelial cells followed PZP cells while some epithelial cells moved as single cells (Suppl. 1G). Given that all KTB epithelial cell lines, independent of ancestry, exhibited strikingly similar leader-follower behaviors during invasion, most of the investigations below describe findings with the KTB34 cell line. In all cases, the phenotypes observed were examined in and extended to all KTB epithelial cell lines except where indicated.

**Figure 1:**
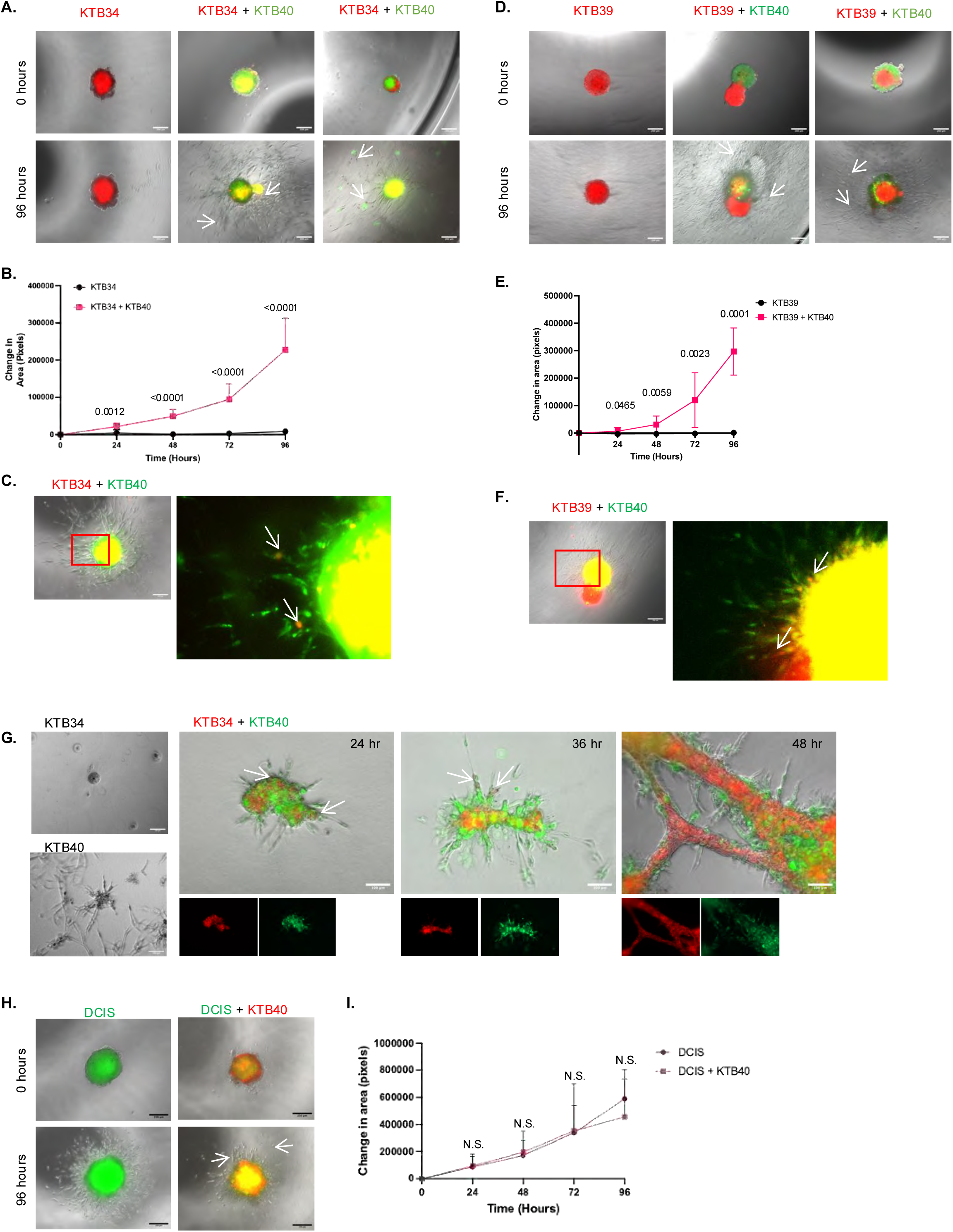
PZP cells confer invasive potential to normal epithelial cells. **A)** Representative images of KTB34 (red) spheroids, and cocultured KTB34 (red) and KTB40 (green) spheroids. Arrows point to invading KTB34 cells in coculture conditions. Scale bar, 200 μm. **B)** Area of epithelial invasion was measured at time points indicated. The data represent 3 independent biological replicates and are presented as means ± SDs. The p value was obtained by unpaired 2-tailed t test for each time point. **C)** In coculture spheroids, KTB40 cells (green) act as leader cells to direct KTB34 cells (red) into the matrix. Arrows point to invading KTB34 cells in magnified view of boxed region. Scale bar, 200 μm. **D)** Representative images of KTB39 (red) spheroids, and cocultured KTB39 (red) and KTB40 (green) spheroids. Arrows point to invading KTB39 cells in magnified view of boxed region. Scale bar, 200 μm. **E)** Area of KTB39 epithelial cell invasion was measured at time points indicated. The data represent 3 independent biological replicates and are presented as means ± SDs. The p value was obtained by unpaired 2-tailed t test for each time point. **F)** In coculture spheroids, KTB40 cells (green) act as leader cells to direct KTB39 cells (red, arrows) into the matrix. Magnified view of boxed region is shown. **G)** Representative images of KTB34 and KTB40 monocultures in Matrigel or cocultures of KTB34 (red) and KTB40 (green) cells in Matrigel, taken at indicated times post seeding are shown. In all coculture conditions KTB40:PZP cells act as leader cells and KTB34 cells (arrows) follow. Scale bar, 100 μm. **I)** Representative images of DCIS (green) spheroids, and cocultured DCIS (green) and KTB40 (red) spheroids. Some DCIS cells (arrows) follow KTB40 cells. Scale bar, 200 μm. **J)** Area of DCIS.com cell invasion was measured at time points indicated. The data represent 3 independent biological replicates and are presented as means ± SDs. The p value was obtained by unpaired 2-tailed t test for each time point.

**Table 1:** Ancestral, phenotypic and molecular characterization of KTB epithelial and stromal cell populations.

| Cell Line | Cell Type | Ancestry Description | Phenotypic and Molecular Characterizations |
| --- | --- | --- | --- |
| KTb8 | Normal ductal epithelium | African Ancestry | Cobblestone morphology |
| KTb34 | Normal ductal epithelium | European Ancestry | Cobblestone morphology, Luminal-A subtype, KRT14+/KRT19+ |
| KTb36 | Normal ductal epithelium | European Ancestry | Cobblestone morphology, Basal subtype |
| KTb39 | Normal ductal epithelium | African Ancestry | Cobblestone morphology, Normal-like subtype, lacks luminal features KRT14+/KRT19+ |
| KTb40 | Fibroadipogenic/mesenchymal stem cell | African Ancestry | Fusiform morphology, PROCR + /ZEB1 + /PDGFRα+, CD105 high/CD26 low, Ecad-/Vim+, transdifferentiate into adipocytes and osteocytes |
| KTb42 | Fibroadipogenic/mesenchymal stem cell | African Ancestry | Fusiform morphology, PROCR + /ZEB1 + /PDGFRα+, CD105 high/CD26 low, Ecad-/Vim+, transdifferentiate into adipocytes and osteocytes |

To complement the spheroid-based assay, we also seeded cells in growth factor reduced Matrigel, a reconstituted basement membrane from Engelbreth-Holm-Swarm (EHS) mouse sarcoma. The KTB epithelial cell lines alone formed compact spheroids 48 hours post seeding, whereas PZP cells formed loose clusters and appeared fibroblast-like. However, when KTB cells were co-cultured with PZP cells, the epithelial cells organized alongside PZP cells in clusters that then developed into long branched tubules as the cells invaded the matrix (Fig. 1G).

To examine if the effect of PZP cells on epithelial cell invasion extended to other epithelial cell types besides the KTB cell lines, we employed the MCF10A cell line which was isolated from an EA patient with fibrocystic breast disease and is commonly recognized as a model for non-transformed mammary ductal epithelium^22,23^. Spheroids composed exclusively of MCF10A cells displayed little invasion into the matrix as expected whereas cocultured MCF10A:PZP spheroids behaved similarly to KTB spheroids, wherein increased epithelial cell invasion and leader-follower behavior was observed (Suppl. 1H, I).

Since PZP cells conferred invasive capacity to normal mammary epithelium, we next asked whether PZP cells induced invasion in a model for preinvasive ductal carcinoma. The MCF10DCIS.com (herein referred to as DCIS.com) cell line is a model of ductal carcinoma in situ that will form comedo DCIS-like lesions *in vivo* and displays a propensity to invade in a collagen matrix *in vitro*^24^. When cultured alone, DCIS.com spheroids invade the collagen matrix (Fig. 1H I). When the DCIS.com cell line was cocultured with PZP cells, we observed some leader-follower behavior, however there was no significant change in extent of DCIS invasion into the matrix in the presence of PZP cells (Fig. 1H, I). Thus, while PZP cells may facilitate and even provide directionality to DCIS movement, they were not required for DCIS cell invasion to the matrix. The results described above also suggest that PZP cells trigger invasive behaviors in cell models of normal ductal epithelium.

### Normal and cancer-associated fibroblasts interact with mammary epithelium differently than PZP cells

Next, we assessed how KTB epithelial cells would respond to other stromal cell populations, fibroblasts in particular. Normal fibroblasts can promote migration and invasion in breast cancer cells however this effect is exacerbated when cancer-associated fibroblasts (CAFs) are cocultured with breast cancer cells^25^. To this end, we cocultured BJ fibroblasts, a normal human foreskin fibroblast line, with KTB34 cells at a 1:1 ratios using our spheroid assays. Coculture of BJ fibroblasts showed modest enhancement of epithelial cell invasion, although the fibroblasts alone invaded the matrix (Fig. 2A-B).

**Figure 2:**
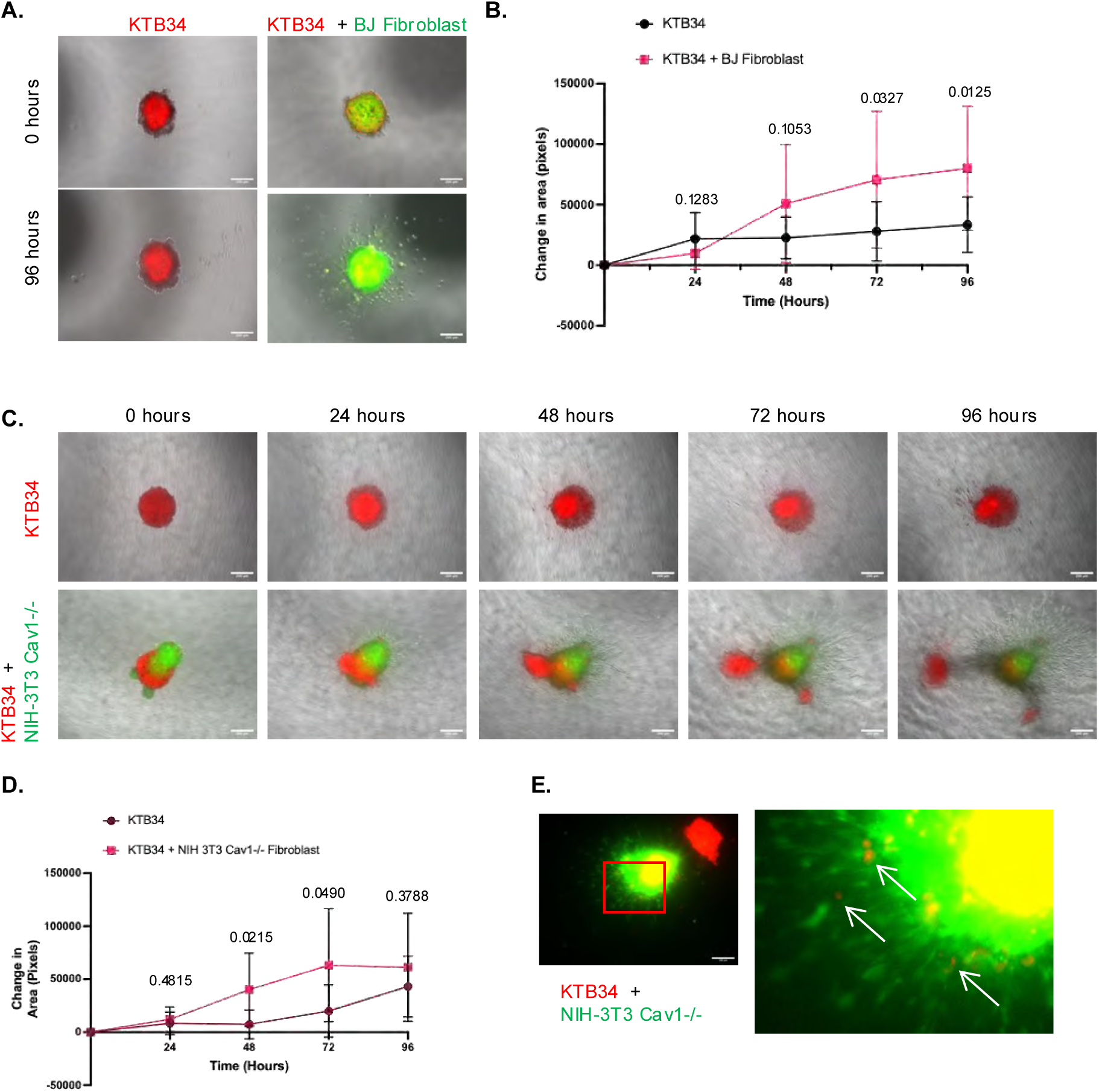
Normal and cancer associated fibroblasts interact with mammary epithelium differently than PZP cells. **A)** Representative images of KTB34 (red) spheroids, and cocultured KTB34 (red) and BJ fibroblasts (green) spheroids. Scale bar, 200 μm. **B)** Area of KTB34 cell invasion was measured at time points indicated. The data represent 3 independent biological replicates and are presented as means ± SDs. The p value was obtained by unpaired 2-tailed t test for each time point. **C)** Representative images of KTB39 (red) spheroids and cocultured KTB39 (red) and BJ fibroblasts (green) spheroids. Scale bar, 200 μm. **D)** Area of KTB34 epithelial cell invasion was measured at time points indicated. The data represent 3 independent biological replicates and are presented as means ± SDs. The p value was obtained by unpaired 2-tailed t test for each time point. **E)** In coculture spheroids, some KTB34 cells (red, arrows) follow NIH 3T3 Cav1-/- cells (green) into the matrix.

We also examined the effects of NIH-3T3 Cav-1 -/- fibroblast line, which displays CAF-like characteristics such as functional inactivation of tumor suppressor RB^26^. In coculture spheroids the KTB34 cells were less intermixed with the CAF cells, forming partial lobes similar to the KTB39 coculture spheroids (Fig. 2C,D). As the NIH-3T3 Cav -/- fibroblasts cells invaded the matrix, some of the KTB34 cells followed invasion into the matrix and the majority of the KTB34 spheroid was separated out from the CAFs at later time points (Fig. 2C, D). Statistically significant invasion occurred by 48 and 72 hour time points, and while some leader-follower invasion was observed, the increase in KTB34 invasion was to a far lesser extent relative to that observed in PZP:epithelial cell coculture spheroids described above (Fig. 2D, F). Thus, PZP cells affect and promote normal epithelial cell invasion more effectively relative to normal fibroblast and CAFs.

### PZP cell secretome activates AKT signaling in epithelial cells that is necessary but not sufficient for epithelial cell invasion

Stromal cells are known to secrete factors to modulate cell behavior in the TME. Previous work demonstrated that coculture of PZP cells with epithelial cells results in the production of secreted factors such as IL-6 and transgelin that can impinge upon signaling cascades involved in invasion^15^. Both IL-6 and transgelin production are known to activate AKT^27,28^ and there is clinicopathologic evidence of significant AKT activation in early breast cancer invasion^29^. This led us to examine whether paracrine signaling from PZP and epithelial cell cocultures increased activation of AKT in KTB epithelial cells. To this end, we treated KTB34 cells with naïve media, KTB34-conditioned media, PZP-conditioned media or KTB34:PZP-coculture media, and then probed for activated/phosphorylated AKT (pAKT). Indeed, treatment of KTB34 cells with PZP cell conditioned media induced an increase in pAKT (Fig. 3A, B). We note that addition of coculture conditioned media increased pAKT abundance in KTB34 cells but to a lesser extent than monoculture PZP media and is likely due to secreted PZP factors being diluted under 1:1 coculture conditions. Interestingly, KTB39 did not display a significant increase in pAKT levels when treated with conditioned media in the same manner compared to KTB34 cells, however, we note that KTB39 cells have a higher baseline level of pAKT (Fig. 3C, D). This suggests that epithelial cells of AA origin maybe primed for processes such as cell invasion.

**Figure 3:**
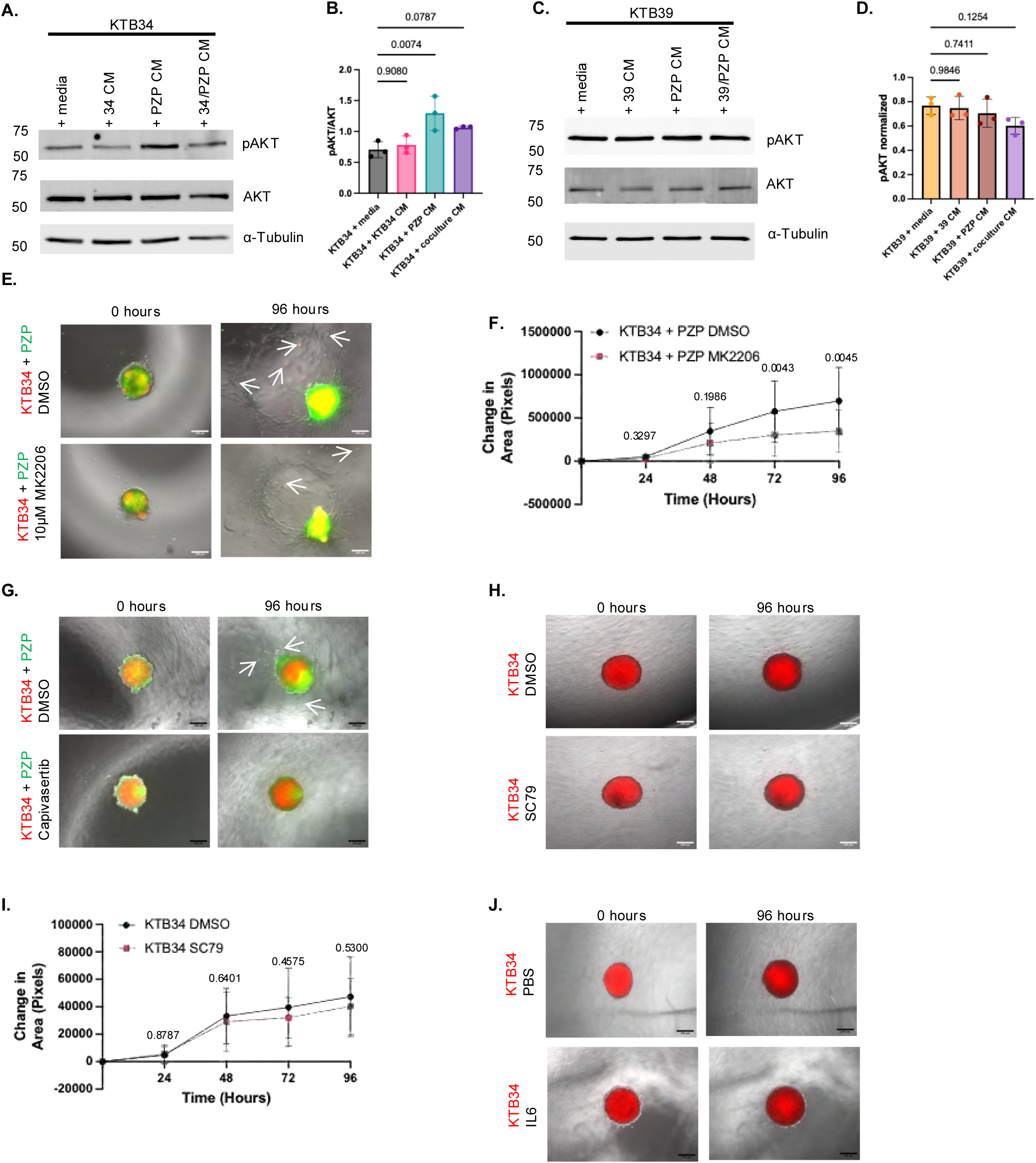
PZP cells impact epithelial cell invasion in part through AKT signaling. **A)** KTB34 cells were treated with naïve media control, KTB34 conditioned media (34 CM), PZP conditioned media (PZP CM) or coculture KTB34:PZP CM (34:PZP CM). Lysates were probed for pAKT (Ser 743), total AKT and α-tubulin (loading control) by western blotting. Images are representative of 3 independent biological replicates. **B)** Densitometry analysis of pAKT levels normalized to total AKT levels. Data are representative of 3 independent biological replicates and are presented as means ± SDs. The p value was obtained by one-way ANOVA with Dunnett’s test for multiple comparisons. **C)** KTB39 cells were treated with naïve media control, KTB39 conditioned media (39 CM), PZP conditioned media (PZP CM) or coculture KTB39:PZP CM (39:PZP CM). Lysates were probe for pAKT (Ser 743), total AKT and α-tubulin (loading control) by Western blotting. Images are representative of 3 independent biological replicates. **D)** Densitometry analysis of pAKT levels normalized to total AKT measured in ImageJ. Data are representative of 3 independent biological replicates and are presented as means ± SDs. The p value was obtained by one-way ANOVA with Dunnett’s test for multiple comparisons at each time point. **E)** Representative images of cocultured KTB34 (red) and PZP (green) spheroids treated with DMSO vehicle control or 10μM MK2206, AKT inhibitor. The data represent 3 independent biological replicates. Arrows point to KTB34 cells. Scale bar, 200 μm. **F)** Area of KTB34 epithelial cell invasion was measured at time points indicated. The data represent 3 independent biological replicates and are presented as means ± SDs. The p value was obtained by unpaired 2-tailed t test for each time point. **G)** Representative images of cocultured KTB34 (red) and PZP (green) spheroids treated with DMSO vehicle control or 10 μM Capivasertib. Arrows point to KTB34 cells. Scale bar, 200 μm. **H)** Representative images of KTB34 (red) spheroids treated with DMSO vehicle control or 20μM SC79, AKT activator. The data represent 3 independent biological replicates. Scale bar, 200 μm. **I)** Area of KTB34 epithelial cell invasion was measured at time points indicated. The data represent 3 independent biological replicates and are presented as means ± SDs. The p value was obtained by unpaired 2-tailed t test for each time point. **J)** Representative images of KTB34 (red) spheroids treated with PBS vehicle control or IL6. Scale bar, 200 μm.

In light of these findings, we investigated the effect of a small molecule inhibitor designed to reduce AKT phosphorylation on spheroid co-cultures. Treatment with 10μM AKT inhibitor^24^ MK2206, significantly reduced pAKT levels (Suppl. 2A-D) and resulted in a marked attenuation in PZP-induced invasion of both KTB34 and KTB39 epithelial cells after 48 hours (Fig. 3E, F, Suppl. 2E, F) indicating that activation of AKT is necessary for epithelial cells to acquire invasive capacity. In addition to abrogation of epithelial cell invasion, pharmacological inhibition of AKT also appeared to reduce PZP cell invasion. Capivasertib, an FDA approved AKT inhibitor^30^ exhibited similar effects on PZP-induced cell invasion (Fig 3G, Suppl. 2G). Since blocking AKT activity resulted in a significant decrease in epithelial cell invasion, we treated KTB34 spheroids with the AKT activator SC79^31^, to determine if AKT activation is sufficient to induce epithelial cell invasion. Treatment with 20 μM SC79 was sufficient to increase pAKT levels in KTB34 cells (Suppl. 2H, I) but did not promote KTB34 epithelial invasion (Fig. 3G, H). Thus, while increased levels of pAKT in epithelial cells is necessary for PZP-induced invasion, AKT activation alone is insufficient to drive this process. This contention is further supported by the data in the KTB39 cell line described above as KTB39 spheroids alone did not invade despite their higher basal levels of pAKT. Furthermore, treatment of KTB spheroids with IL-6 alone did not support epithelial cell invasion however it promoted AKT activation, (Fig 3J, Suppl. 2J). These findings suggest that either additional PZP-secreted factors required for invasion were diluted out in the co-culture media, and/or that the physical presence of the PZP cells facilitates KTB epithelial cell invasion.

### Characterization of PZP cell-facilitated epithelial cell invasion

Epithelial cells can undergo partial epithelial-to-mesenchymal transition (EMT) to become more invasive by down regulating E-cadherin and increasing N-cadherin expression^32–34^. We investigated if the PZP stimulated epithelial cells for epithelial-to-mesenchymal transition (EMT) markers. KTB34 cells were treated with either naïve media control, KTB34 conditioned media, PZP conditioned media or coculture KTB34:PZP conditioned media and then probed for the classical EMT markers E-cadherin, N-cadherin and vimentin. We observed no change in levels or distribution of these markers indicating that the epithelial cells maintain their epithelial identity despite the increased invasive behavior (Fig. 4A-D). Similarly treated epithelial cell spheroids exhibited no cell invasion in any experimental condition indicating that conditioned media alone is insufficient to induce invasion (Fig. 4E). Because AKT activation also plays a significant role in breast cell proliferation^35^, we treated KTB34 cells with the same conditioned media and examined proliferation over 48 hours. Crystal violet staining demonstrated no change in cell proliferation (Fig. 4F). Thus, while PZP cell conditioned media induced the necessary activation of AKT, there was no effect on EMT status or cell proliferation.

**Figure 4:**
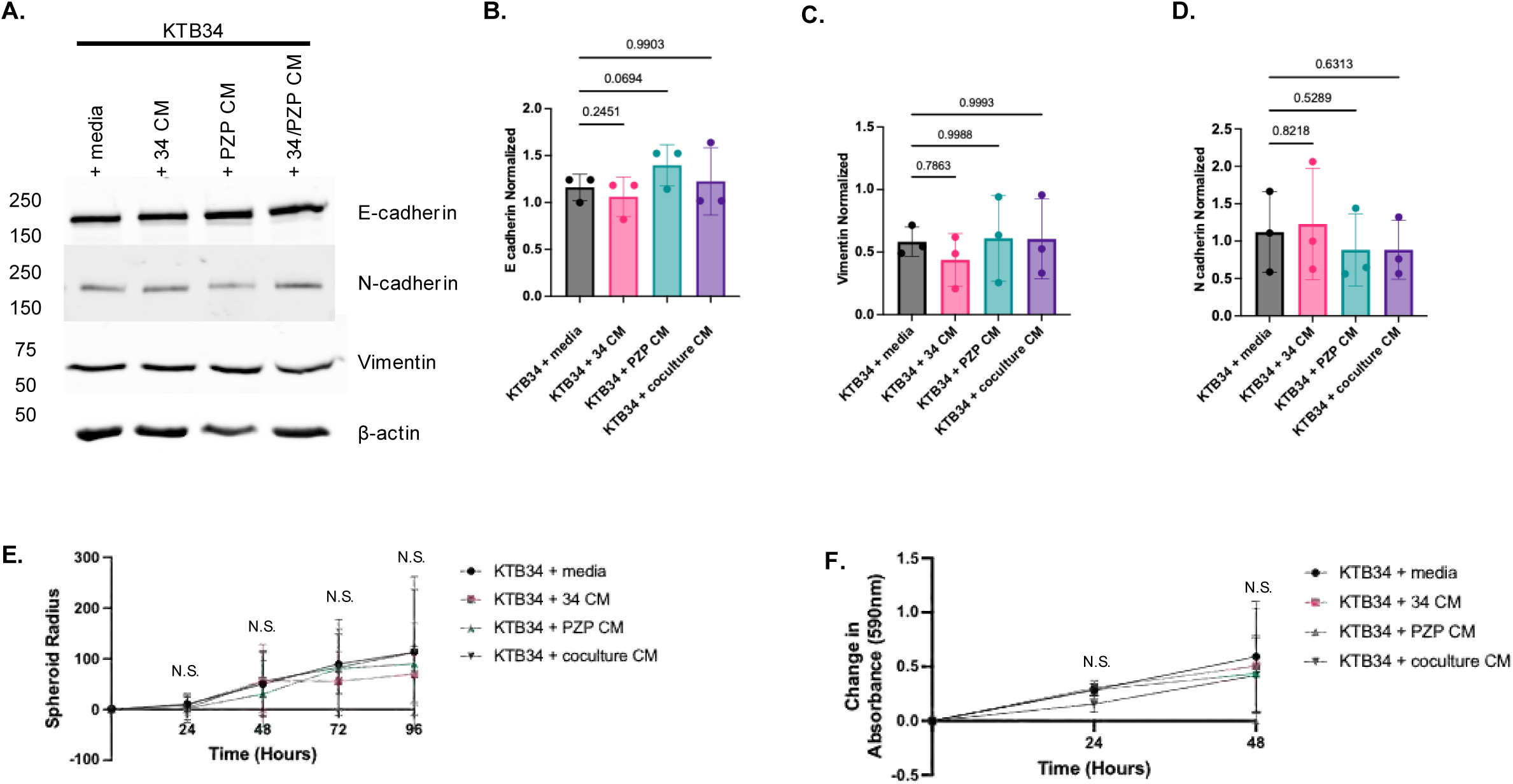
PZP cell secretome does not affect EMT markers, epithelial cell invasion, or proliferation. **A)** KTB34 cells were treated with naïve media control, KTB34 conditioned media (34 CM), PZP conditioned media (PZP CM) or coculture KTB34:PZP CM (34:PZP CM). Lysates were probed for classical EMT markers E-cadherin, N-cadherin, Vimentin and β-actin (loading control) by Western blotting. Images are representative of 3 independent biological replicates. **B)** Densitometry analysis of E-cadherin levels normalized to β-actin. Data are representative of 3 independent biological replicates and are presented as means ± SDs. The p value was obtained by one-way ANOVA with Dunnett’s test for multiple comparisons. **C)** Densitometry analysis of vimentin normalized to β-actin. Data are representative of 3 independent biological replicates and are presented as means ± SDs. The p value was obtained by one-way ANOVA with Dunnett’s test for multiple comparisons. **D)** Densitometry analysis of N-cadherin abundance normalized to β-actin abundance measured in ImageJ. Data are representative of 3 independent biological replicates and are presented as means ± SDs. The p value was obtained by one-way ANOVA with Dunnett’s test for multiple comparisons. **E)** Monoculture KTB34 spheroids were subjected to treatment with naïve media control, KTB34 conditioned media (34 CM), PZP conditioned media (PZP CM) or coculture KTB34:PZP CM (34:PZP CM). Spheroid radius was measured by taking cross-sectional measurements of furthest point to the center of the spheroid in ImageJ. The data represent 3 independent biological replicates and are presented as means ± SDs. The p value was obtained by one-way ANOVA with Dunnett’s test for multiple comparisons at each time point. **F)** KTB34 cells were subjected to treatment with naïve media control, KTB34 conditioned media (34 CM), PZP conditioned media (PZP CM) or coculture KTB34:PZP CM (34:PZP CM) at a ratio of 2:1 with complete media. Cells were fixed with 1% glutaraldehyde and stained with 0.1% crystal violet. Stained samples were solubilized with 0.2% and measured at 590nm wavelength. Data are representative of 3 independent biological replicates and are presented as means ± SDs. The p value was obtained by one-way ANOVA with Dunnett’s test for multiple comparisons at each time point.

### PZP cells influence ECM to guide mammary ductal epithelium

Stromal cell populations have been shown to deposit ECM proteins to facilitate invasion^36,37^. Western blot analysis demonstrated that PZP cells alone produce more fibronectin (which is known to bind β1-integrin) than KTB34 cells (Fig. 5A, B). Notably however, there was a further increase in the amount of fibronectin when cells were cocultured (Fig. 5A, B). To assess fibronectin deposition during invasion, we used CellTracker dye to stain the KTB34 population of cells and monitored spheroid formation in 1:1 coculture with unstained PZP cells. The collagen matrices were partially digested with collagenase and stained for fibronectin and β1-integrin. Fibronectin appeared closely aligned alongside the integrin-labeled invasive cells at the edge of the spheroid, and adjacent to blue dyed KTB34 cells (Fig. 5C). We note that while both KTB34 and PZP cells express β1-integrin, PZP cells express higher levels (Suppl. 2K) allowing visualization of the higher signal in PZP cells by immunofluorescence microscopy. Further analysis by immunofluorescence in 2D shows fibronectin deposits exclusively by PZP cells and a network of fibronectin in cocultured cells (Fig. 5D). Similar to western blotting data, fluorescent intensity quantification also showed that there was an increase in deposited fibronectin when PZP cells were cocultured with KTB34 cells compared to the individual cultures, raising the possibility of paracrine signaling from epithelial cells (Fig. 5E).

**Figure 5:**
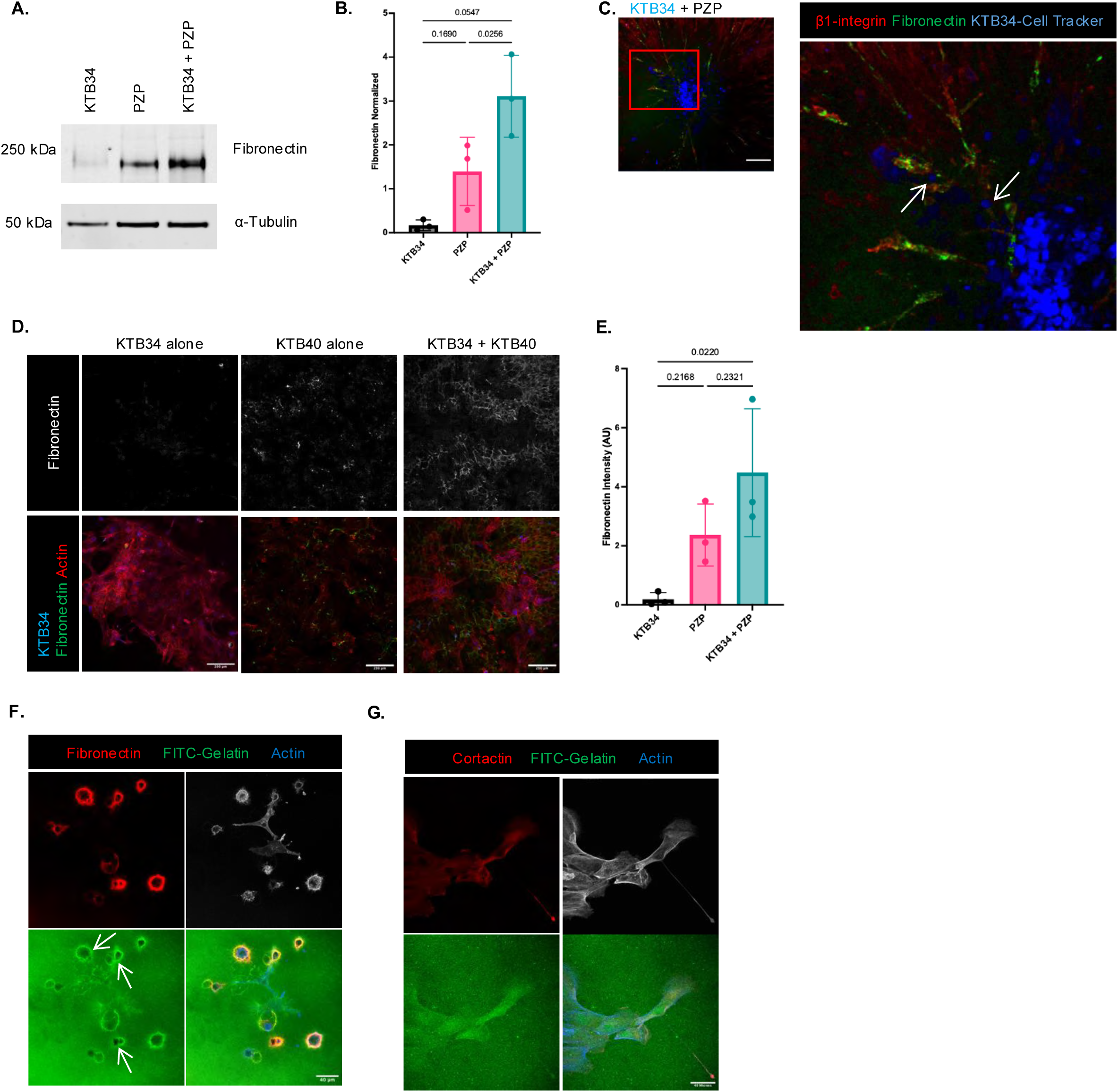
PZP cells influence ECM to allow epithelial cell invasion. **A)** Coculture of KTB34 with PZP cells increases fibronectin abundance. KTB34, PZP and cocultured KTB34 and PZP whole cell lysates were probed or fibronectin and for α-tubulin as a loading control, by Western blotting. Images are representative of 3 independent biological replicates. **B)** Densitometry analysis of fibronectin levels normalized to α-tubulin measured in ImageJ. Data are representative of 3 independent biological replicates and are presented as means ± SDs. The p value was obtained by one-way ANOVA with Tukey’s correction for multiple comparisons. **C)** Representative spheroids with immunofluorescent staining of KTB34 (Cell Tracker Dye, blue) cocultured with PZP cells followed by staining for β1-integrin (red) and fibronectin (green). Images shown as max projection. KTB34 cells (arrow) invade into the matrix following β1-integrin positive cells (PZP) that produce fibronectin. Scale bar, 200 μm. **D)** KTB34 (blue), PZP and coculture of KTB34 (blue) and PZP cells were fixed, stained as indicated, and imaged by confocal microscopy. Fibronectin is primarily produced by PZP cells. Images are representative of 3 biological replicates. **E)** Quantification of fibronectin fluorescence intensity as described in D. Data are presented as means ± SDs. The p value was obtained by one-way ANOVA with Tukey’s correction for multiple comparisons. **F)** Cocultures of KTB34 and PZP cells were subjected to the scratch assay, fixed at times indicated, and labeled for E cadherin (blue), fibronectin (green), and β1-integrin (red). Highly positive β1-integrin cells (PZP cells, marked by asterisks) deposit fibronectin while KTB34 cells retain E-cadherin labeling. Scratch denoted by yellow line. Scale bar, 100 μm. **G)** PZP cells on FITC-labeled gelatin were labeled for fibronectin (red) and actin (blue). PZP cells degrade the gelatin and fibronectin deposits around areas of matrix degradation (arrows) are shown. Images are representative of 3 independent biological replicates. **H)** KTB34 cells were seeded on FITC labeled gelatin and labeled for cortactin (red) and actin (blue). Little to no gelatin degradation was observed.

Previous studies have shown that fibroblast-led micro tracks allow invasion in a dense collagen environment in an MMP-dependent manner^38^. To investigate if this was the case for PZP cells, we employed the gelatin invasion assay to determine if the PZP cells were capable of degrading the extracellular matrix to provide a path for invasion^39^. KTB40 cells seeded on thin FITC-gelatin degraded the matrix below the cells and deposited fibronectin in these areas of degradation (Fig. 5G). In contrast, under the same experimental conditions KTB epithelial cells show little to no invasion of gelatin (Fig. 5H). These data collectively suggest that PZP cells create paths of least resistance through matrix deposition and further, PZP-coupled fibronectin deposition may provide guidance cues facilitating directed movement of epithelial cells.

## Discussion

There is much that remains undiscovered about the racial and ethnic disparities in breast cancer incidence and mortality rates, but we do know that the risk factors include various socioeconomic factors as well as the biological differences between racial/ethnic populations^7^. Thus, to improve outcomes for AA breast cancer patients, who have an estimated 41 percent higher death rate from breast cancer compared to other populations, it is important to address both socioeconomic disparities as well as further characterize the biological basis for the increased onset and progression seen among AA patient populations. In this study, we have investigated the contributions of a relatively newly described stromal cell type, referred to as PZP cells, on the acquisition of invasive potential in human breast epithelial cells. PZP cells, characterized by their expression of <u>P</u>ROCR, <u>Z</u>EB1 and <u>P</u>DGFRα, are present at intrinsically higher levels in the normal breast tissue from AA donors compared to donors of any other ancestry^14,15^. Moreover, the numbers of PZP cells are increased in breast cancer tissues in EA patients, suggesting that these cells play a significant role in breast cancer progression.

We report here that PZP cells stimulate the invasive capacity of human breast epithelial cells in the following ways. First, the PZP cells promote the activation of AKT in epithelial cells and this activation is necessary for epithelial cell invasion. Notably, KTB39 cells, which are of African ancestry, displayed higher baseline levels of pAKT compared to the KTB34 cell line of EA origin. Interestingly in this regard, KTB39 is unique in that it lacks luminal epithelial characteristics^14^. Blocking AKT activation significantly reduced epithelial cell invasion in the 3D spheroid coculture model regardless of genetic ancestry, indicating this is an important signaling node in epithelia for PZP cell-mediated invasion. Second, PZP and epithelial cell interactions resulted in a significant increase in the production and deposition of the ECM protein, fibronectin, which likely provides guidance cues to facilitate directed cell movement. Finally, PZP cells promoted the proteolytic breakdown of the ECM creating paths of least resistance during cell invasion. These findings support the contention that intrinsically higher numbers of PZP cells could predispose pre-cancerous cells in AA tissues to become invasive and aggressive during disease progression. EA tissues on the other hand, may require additional physiological processes to promote expansion of PZP cells. We note however, that we have not investigated PZP cells derived from EA tissues in this study.

Previous work has demonstrated that coculture of PZP cells with epithelial cells results in the production of secreted factors, such as IL-6 and transgelin, both of which are known to activate AKT^27,28^. Indeed in our system, addition of IL-6 results in the activation of AKT in the KTB34 epithelial cell line. The PZP secretome alone could promote AKT activation in epithelial cells supporting the contention that paracrine signaling from PZP cells facilitates AKT activation in epithelial cells. However, activation of AKT alone is insufficient to induce epithelial cell invasion. As stated above, one of the two AA cell lines examined, KTB39 cells, displayed higher baseline levels of pAKT compared to the other KTB epithelial cell lines. We speculate, and it remains to be determined, if this elevated level of activated AKT intrinsically primes AA epithelial tissues for acquisition of invasive properties.

PZP cells allow epithelial cell invasion to a higher degree than normal fibroblasts and CAFs based on the spheroid invasion assays. CAFs that induce invasion in malignant breast cancer cells promote EMT in the epithelial cells^40–42^. We did not observe a change in epithelial status in KTB34 cells when cocultured or treated with CM from the PZP cells. PZP cells share characteristics of fibroblasts but also can transdifferentiate into adipocytes^15^. Like fibroblasts, adipose stromal cells have also been shown to stiffen the tumor microenvironment by depositing collagen and fibronectin^43^. High fibronectin expression and tumor stiffness are associated with decreased overall and disease-free survival in the clinic^44^.

PZP cells when cocultured with the tumorigenic DCIS.com cell line displayed leader-follower behavior to facilitate invasion but did not increase the invasiveness of the DCIS.com cells. Thus, it appears that PZP cells may have most impact on normal epithelial cells, and therefore at very early stages of disease, although PZP cells could potentially also affect other aspects/stages of tumorigenesis. Patients of African descent are more frequently diagnosed with TNBC, which is one of the most aggressive types of breast cancer and has more limited options for treatment^45,46^. Characterization of the PZP cells points to potential clinical targets, expanding options for patients. Notably, PDGFR, a marker of PZP cells, is a druggable target and stromal PDGFα activation increases mammary matrix stiffness and tumor growth^47–49^. Recent clinical trials have investigated the effectiveness of AKT inhibitors in treating advanced stage and metastatic breast cancer^50,51^. Capivasertib, an AKT inhibitor has received approval by Federal Drug Administration approved for hormone receptor positive, HER2^-^ breast cancer with biomarker alterations in the PI3K/AKT pathway^51^. In our 3D coculture spheroid model, we found Capivasertib effectively blocked invasion by epithelial and PZP cells. Targeting the PZP cell population and pathways influenced by PZP cells at earlier time points may provide a complementary or alternative avenue for treating more aggressive tumor types.

In summary, this study shows that PZP cells influence epithelial cell behavior through a combination of activating AKT signaling and through the physical remodeling of the extracellular matrix by depositing fibronectin. The presence of the PZP cell lines induces invasive behavior in epithelial cells regardless of genetic ancestry. Together these data reveal multiple mechanisms in which PZP cells may influence breast tumorigenesis and furthers knowledge of the molecular basis underpinning disparate patient prognosis and outcomes in populations of African ancestry.

## Acknowledgements

We thank Brijesh Kumar from the Nakshatri lab for his guidance with growing and maintaining KTB cell lines, and Sara Cole from the Notre Dame Integrated Imaging Facility for her assistance with confocal microscopy and utilization of imaging software. This work was supported in part by NIH/NCI CA273469 to CDS and a grant from the 100 Voices of Hope to HN and CDS.

## Author Contributions

MGS: Developed the study, formulated the methods, conducted experiments, collected and analyzed data, wrote the initial draft, edited the manuscript and prepared figures. VE: Assisted with preparing reagents and conducted experiments. JC: conducted experiments, collected and analyzed data, provided supervision. HN: sharing of KTB cell lines, data analysis, review of manuscript. CDS: Responsible for overall project management including conceptualization, project development, assisting with experimental design, analyzing data and editing the manuscript.

## MATERIALS AND METHODS

### Cell Culture

KTB34, KTB36, KTB39, KTB8, KTB40 and KTB42 cell lines obtained from the Nakshatri lab, have been described^14,15,52^. KTB cells were maintained in 1:3 DMEM-low glucose (Gibco) with 25 mM HEPES (Sigma-Aldrich) and F-12 nutrient (Gibco) supplemented with 10% FBS (Atlas),

0.4 μg/ml hydrocortisone (Sigma-Aldrich), 1% penicillin-streptomycin solution (Gibco), 5 μg/ml Insulin (Sigma-Aldrich), 20 ng/ml EGF (R&D Systems), 24 μg/ml Adenine (Sigma-Aldrich), 100 nM ROCK inhibitor (Enzo Life Sciences). BJ fibroblasts were maintained in EMEM supplemented with 10% FBS (Atlas), 1mM sodium pyruvate (Gibco), 1X L-glutamine, and 1% penicillin-streptomycin solution (Gibco). NIH-3T3 Cav-1 -/- fibroblasts were kindly provided by Dr. Zachary Schafer, University of Notre Dame, and were maintained in DMEM (Gibco) supplemented with 10% FBS and 1% penicillin-streptomycin solution (Gibco) as described^53^. MCF10DCS.com cells^54^ were maintained in DMEM/F-12 supplemented with 10% FBS, 2 mM L-glutamine, and 1% penicillin-streptomycin solution (Gibco). MCF-10A cells were kindly provided by Dr. Joan Brugge, Harvard University School of Medicine and maintained in DMEM/F12 supplemented with 5% horse serum, 2 mM L-glutamine, 20 ng/mL EGF, 500 ng/mL hydrocortisone, 100 ng/mL cholera toxin, 10 mg/mL insulin, and 100 U/mL penicillin-streptomycin as described^55^. All cell lines were incubated at 37°C, with 5% CO_2_ and were routinely tested for *Mycoplasma*. To generate conditioned media, active log-phase cultures of all cells were trypsinized, resuspended in growth media and counted. An equal total number of 200,000 cells were plated individually and in coculture in a 6 well plate. Cells were allowed to reach confluence and then media was replaced with serum free culture media with no additives. After 24 hours, conditioned media was collected and stored at 4°C if not immediately used.

### Antibodies

Primary antibodies utilized were anti-AKT (Cell Signaling Technologies, 9272), anti-phospho-AKT S473 (ProteinTech, 80455-1-RR), anti-alpha-tubulin (ProteinTech, 66031-1-Ig), anti-fibronectin (ProteinTech, 15613-1-AP), anti-E-cadherin (BD Transduction Labs, 610181), anti-N-cadherin (Cell Signaling, 14215P), anti-Vimentin (ProteinTech, 60330-I-Ig), anti-β-1 integrin (DSHB, AIIB2), anti-cortactin (EMD Millipore, 05-180). Alexa Fluor Plus 647 Phalloidin was from Invitrogen, A30107. Secondary antibodies used were Donkey anti-mouse Alexa Fluor Plus 555 (ThermoFisher, catalog # A32773), Donkey anti-mouse Alexa Fluor Plus 680 (ThermoFisher, catalog # A32766), Donkey anti-rabbit Alexa Fluor Plus 488 (ThermoFisher, catalog # A32790), Donkey anti-mouse Alexa Fluor Plus 555 (ThermoFisher, catalog # A32773), Donkey anti-rabbit Alexa Fluor Plus 800 (ThermoFisher, catalog # A32808), and Donkey anti-Rat Alexa Fluor Plus 647 (ThermoFisher, catalog # A48272).

### Spheroid culture

Active log-phase cultures of all cell types were trypsinized, resuspended in growth media and counted. 10,000 cells per well were seeded in ultra-low attachment U-bottom 96 well plates and incubated in growth media for 3 days to form spheroids. For spheroid embedment, a solution of 10% PureCol (Advanced BioMatrix) with 10X PBS was prepared and pH adjusted to approximately 7.0. A layer of PureCol/PBS solution was added to a flat bottom 96 well plate and allowed to solidify. Spheroids were individually harvested and transferred in a 10% PureCol with 10X serum free DMEM (pH approximately 7.0) solution to individual wells for embedment. The PureCol/10x DMEM solution was allowed to solidify before adding complete growth media. Spheroids were imaged every 24 hours with a Zeiss Axio Observer. For spheroid invasion quantification, the area of invasion was measured in ImageJ based on total area occupied by epithelial cells at each time point. Images were split into individual fluorophore channels and the brightness in the channel corresponding to the epithelial cells was adjusted to allow invading cells to be visualized. Area of invasion was measured by tracing the core spheroid and radial outlines of invading cells. Individual invasion area was normalized by subtracting the starting area from the area of each time point. For epithelial follower versus single cell quantification, epithelial cells moved out into the matrix that did not appear to be in contact with PZP cells were counted as single cells. Follower cells were defined as epithelial cells in contact and behind PZP cells that invaded into the matrix. Images were split into individual fluorophore channels and the brightness in each channel was adjusted to allow invading cells to be visualized. Data was normalized by calculating percent of cells per phenotype.

### Matrigel assay

For Matrigel cultures, a base layer of thawed growth factor-reduced Matrigel was prepared and allowed to solidify for approximately 90 minutes. Active log-phase cultures of KTB34 and KTB40 cells were trypsinized, resuspended in growth media and counted. Cells were diluted in assay media consisting of culture media with additional 2% Matrigel and 20ng/ml EGF, then seeded on top of the base layer. Cultures were imaged every 24 hours with a Zeiss Axio Observer.

### Immunofluorescence microscopy

Active log-phase cultures of all cell types were trypsinized, resuspended in growth media and counted. An equal total number of 100,000 cells were plated individually and in coculture on cover glass and incubated overnight. Cells were fixed with 2% paraformaldehyde, then washed in 1X PBS with 100 mM glycine (VWR) and incubated in blocking solution composed of 5% BSA, 0.2% TritonX-100, 0.05% Tween-20 (all purchased from VWR) in 1X PBS for 25 minutes. Primary antibodies were incubated at room temperature for 2 hours and washed with 1X IF wash buffer (10X stock: 1.3M NaCl, 132μM NaH_2_PO_4,_ 33.3μM NaH_2_PO_4_, 76.9 μM NaH_3_, 5% BSA, 2% TritonX-100, 0.4% Tween-20 in water, pH 7.4) before incubation with secondary antibodies for 1 hour. Cover glass was mounted with Prolong Gold Antifade mounting media and imaged with a Leica Stellaris 8 DIVE multiphoton microscope housed in the Notre Dame Integrated Imaging Facility, University of Notre Dame. Ten fields per coverslip were chosen at random without bias for each sample. Spheroid cultures were prepared as above and after 48 hours, collagen was partially digested with 100U/mL collagenase for 5 minutes. Spheroids were then fixed with 4% paraformaldehyde for 40 minutes, washed in 1X PBS with 100 mM glycine (VWR), and incubated with blocking buffer for 40 minutes. Primary antibody was incubated at room temperature for 2 hours and spheroids were washed with 1X IF wash buffer before incubating with secondary antibodies for 2 hours. Chambers were removed and then cover glass mounted with Prolong Gold Antifade mounting media. Spheroids were imaged with a Leica Stellaris 8 DIVE multiphoton microscope. For Cell-tracking studies, cells were fluorescently labeled by pre-incubating them with 10 μM CellTracker Green CMFDA (Thermo Fisher, catalog # C7025) or CellTracker Orange CMRA (Thermo Fisher, catalog # C34551) in serum free media for one hour at 37°C with 5% CO_2_.

### Gelatin invasion assay

Thin, gelatin coated cover slips were prepared as previously described^39,56^. Briefly, FITC conjugated gelatin was warmed to 37°C, used to coat flame sterilized cover glass and cross-linked with 1% glutaraldehyde in 1X PBS. Active log-phase cultures of KTB34 or KTB40 cells were trypsinized and counted. 30,000 cells were plated on gelatin coverslips and cells were allowed to invade for approximately 16 hours. Coverslips were prepared as described above and imaged with a BioRad MRC-1024 laser scanning confocal microscope or Leica Stellaris 8 DIVE multiphoton microscope.

### Western blotting

For western blotting, cells were lysed in buffer containing 150 mM NaCl, 25 mM Tris-HCl pH7.6, 1% sodium deoxycholate, 0.1% SDS, and 1% Triton X-100 (all purchased from VWR) with mammalian protease inhibitor cocktail and phosphatase inhibitor cocktail (Millipore Sigma) added to complete the buffer just before use. Equal amounts of protein were mixed with 4X SDS loading dye and incubated in a boiling water bath for 5 minutes. Lysates were then separated by SDS-PAGE and then transferred to PDVF-FL membrane (Millipore Sigma). Non-specific binding was blocked by incubation in 5% nonfat milk in TBS for 1 hour at room temperature. Primary antibody incubation was carried out in 5% nonfat milk in TBS+0.1% Tween-20 overnight at 4°C with gentle rocking. Membranes were washed in TBS+0.1% Tween-20 before incubation with Alexa Fluor Plus secondary antibodies in 5% nonfat milk in TBS for 1 hour at room temperature. Membranes were again washed in TBS+0.1% before being imaged using a LiCor Odyssey scanner.

### Cell proliferation assay

Active log-phase cultures of KTB34 cells were trypsinized, resuspended in growth media and counted. 75,000 cells were seeded per well in 12-well plates. The following day the cells were incubated in serum free media for 3 hours prior to adding 1:2 complete media:conditioned media. Naïve serum free media with no additives used as control media was substituted for conditioned media. Cells were fixed with 1% glutaraldehyde in 1X PBS for 15 minutes at room temperature before adding 0.1% crystal violet in 1X PBS for 30 minutes at room temperature. Excess crystal violet was removed, and plates allowed to dry before solubilizing crystal violet stain in 0.2% TritonX-100 in 1X PBS for 30 minutes. Solubilized crystal violet was measured at 590 nm using a plate reader.

### Statistical Analysis

Relevant statistical details are included in individual figure legends. Statistical analyses were performed using GraphPad Prism 9. Student’s t test was used when comparing two groups with data that appeared to be normally distributed with similar variances. When comparing multiple treatment groups to a single control and the data were normally distributed, we employed one-way ANOVA with Dunnett’s correction for multiple comparisons. When multiple groups were being analyzed and each group was compared to all other groups and the data appeared normally distributed, we performed one-way ANOVA with the Tukey test to correct for multiple comparisons.

## Supplemental Information

**Supplemental Figure 1:**
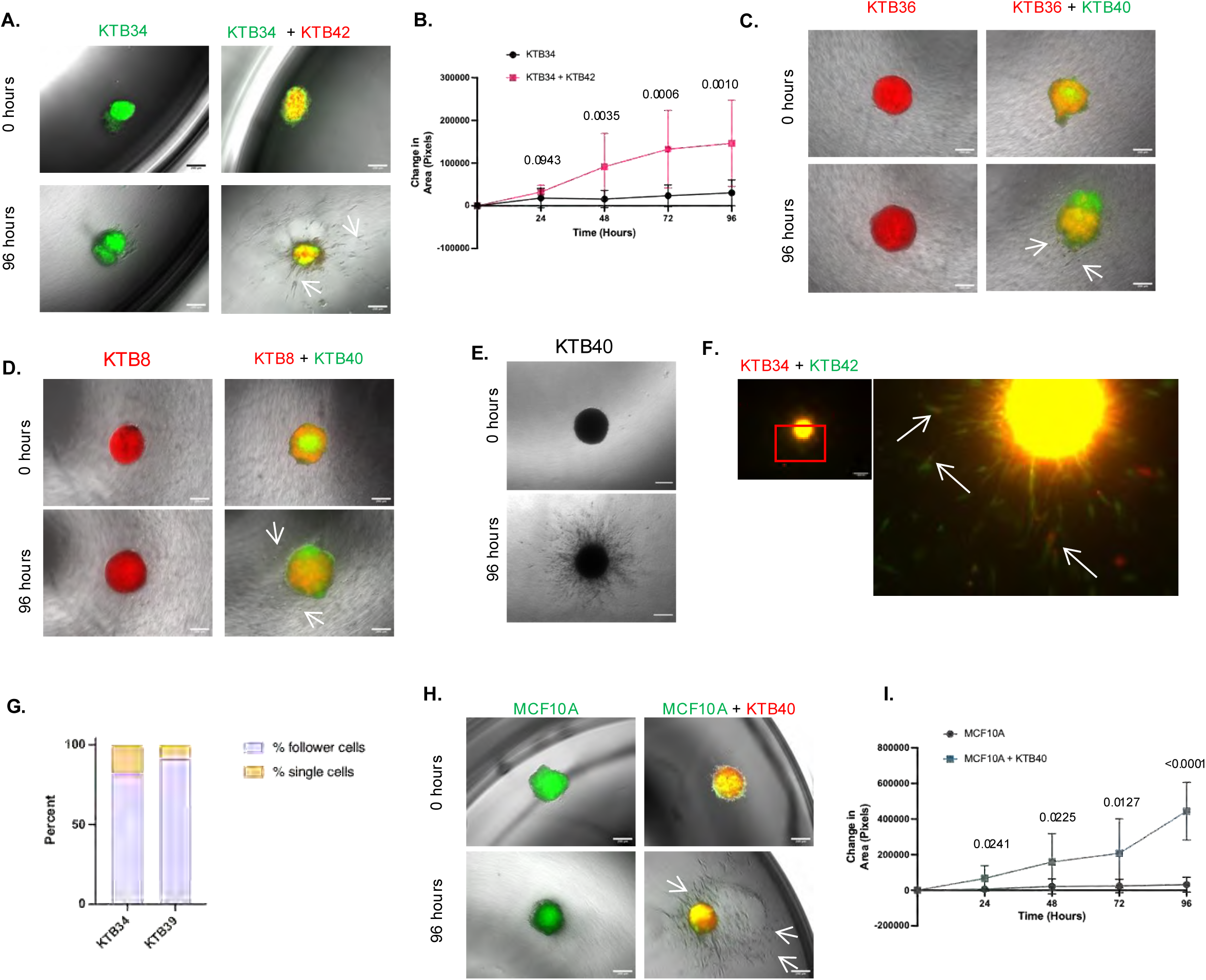
Effect of PZP cells on the invasive potential of normal epithelial cells. **A)** Representative images of KTB34 spheroids, and cocultured KTB34 (green) and KTB42 (red) spheroids. Arrows point to invading KTB34 cells. Scale bar, 200 μm. **B)** Area of KTB34 epithelial cell invasion was measured at time points indicated. The data represent 3 independent biological replicates and are presented as means ± SDs. The p value was obtained by unpaired 2-tailed t test for each time point. **C)** Representative images of KTB36 (red) and KTB36 (red) cocultured with KTB40 (green) spheroids. Scale bar, 200 μm. **D)** Representative images of KTB8 (red) and KTB8 (red) cocultured with KTB40 (green) spheroids. Arrows point to invading KTB8 cells. Scale bar, 200 μm. **E)** Representative images of monoculture KTB40 spheroids. Scale bar, 200 μm. **F)** In coculture spheroids, KTB42 cells (green) act as leader cells (arrows) to direct KTB34 cells (red) into the matrix.. Arrows point to invading KTB34 cells in magnified view of boxed region. **G)** Quantification of the percent of KTB34 or KTB39 cells observed invaded into the matrix and follower or single cells at 48 hours **H)** Representative images of MCF10A (green) and MCF10A (green) cocultured with KTB40 (red) spheroids. Arrows indicate invading MCF-10A cells. Scale bar, 200 μm. **I)** Area of MCF 10A epithelial cell invasion was measured at time points indicated. The data represent 3 independent biological replicates. The p value was obtained by unpaired 2-tailed t test for each time point.

**Supplemental Figure 2:**
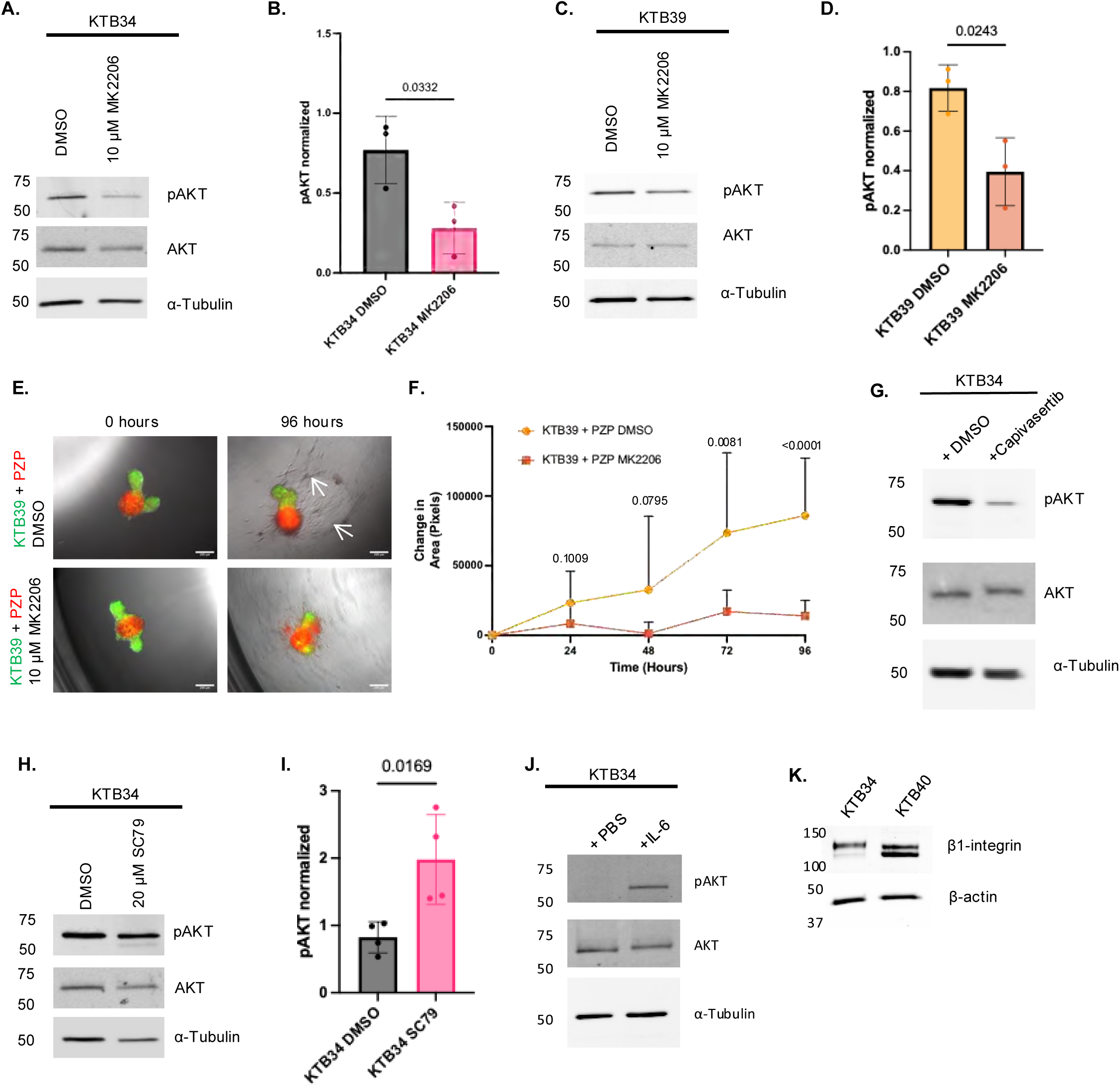
Effect of PZP cells on AKT signaling. **A)** KTB34 cells were treated with DMSO vehicle control or 10 μM MK2206 and probed for pAKT (Ser 473), total AKT and α-tubulin (loading control) by Western blotting. The data represent 3 independent biological replicates. **B)** Densitometry analysis of pAKT levels normalized to total AKT levels. Data are representative of 3 independent biological replicates. The p value was obtained by unpaired 2-tailed t test. **C)** KTB39 cells were treated with DMSO vehicle control or 10 μM MK2206. Lysates probed for pAKT (Ser 473), total AKT and α-Tubulin (loading control) by Western blotting. The data represent 3 independent biological replicates. **D)** Densitometry analysis of pAKT levels normalized to total AKT levels. Data are representative of 3 independent biological replicates and are presented as means ± SDs. The p value was obtained by unpaired 2-tailed t test. **E)** Representative images of KTB39 (green) cocultured with PZP cells (red) spheroids DMSO control or treated with 10 μM MK2206. Arrows point to invading KTB 39 cells. Scale bar, 200 μm. **F)** Area of KTB39 epithelial cell invasion was measured at time points indicated. The data represent 3 independent biological replicates and are presented as means ± SDs. The p value was obtained by unpaired 2-tailed t test for each time point. **G)** KTB34 cells treated with DMSO vehicle control or 10 μM Capivasertib. Lysates were probed for pAKT (Ser 473), total AKT and α-tubulin (loading control) by Western blotting. **H)** KTB34 cells treated with DMSO vehicle control or 20 μM SC79 (AKT activator). Lysates were probed for pAKT (Ser 473), total AKT and α-Tubulin (loading control) by Western blotting. The data represent 3 independent biological replicates. **I)** Densitometry analysis of pAKT normalized to total AKT. Data are representative of 3 independent biological replicates and are presented as means ± SDs. The p value was obtained by unpaired 2-tailed t test. **J)** KTB34 treated with PBS vehicle control or IL6. Lysates were probed for pSTAT3 (Tyr 705), total STAT3 and α-tubulin (loading control). **K)** KTB34 and KTB40 lysates were prepared and probed for β1-integrin and β-actin (loading control) by Western blotting.

